# Choroid plexus enlargement in acute neuroinflammation is tightly interrelated to the tyrosine receptor signalling

**DOI:** 10.1101/2024.03.09.583615

**Authors:** Felix Luessi, Julia Schiffer, Gabriel Gonzalez-Escamilla, Vinzenz Fleischer, Sinah Engel, Dumitru Ciolac, Thomas Koeck, Philipp S. Wild, Joel Gruchot, Tobias Ruck, Ahmed Othmann, Stefan Bittner, Sven G. Meuth, Frauke Zipp, Olaf Stüve, Sergiu Groppa

## Abstract

The choroid plexus (ChP) plays a crucial function in neuroinflammation of the central nervous system and in the immune response of the brain during neurodegeneration. Recent studies described a massive ChP enlargement in patients with multiple sclerosis (MS) and active disease courses, but also in several other neuroinflammatory and neurodegenerative conditions. Nevertheless, the exact basis and pathophysiology behind ChP hypertrophy remains unclear. This study was designed to evaluate the association of cerebrospinal fluid (CSF) proteomic spectra with brain MRI-derived volumetric measures of ChP in two independent cohorts of MS patients, and to translationally validate the related molecular mechanisms in the transcriptomic analysis of the ChP properties in a mouse model of experimental autoimmune encephalomyelitis (EAE). Our analysis revealed five enriched proteins *(NTRK2, ADAM23, SCARB2, CPM, CNTN5)* significantly associated with the ChP volumes in both of the MS cohorts. These proteins relate closely to mechanisms of cellular communication, function (e.g. transmembrane tyrosine receptor signalling (RTK) and vascular endothelial growth) and pathways involved in the regulation of cellular plasticity (e.g. neuron differentiation, axonal remodelling and myelin regulation) as depicted by molecular function analysis and validation of the results in the transcriptome from ChP tissue specific for EAE. This work provides conclusive new evidence for the role of ChP in the context of neuroinflammation and neurodegeneration, demonstrating the intriguing relationships between ChP enlargement, CSF dynamics, and the development of neuroinflammatory and neurodegenerative diseases. Our results are encouraging for the development of new therapeutic avenues (i.e. targeting RTK signalling).

**One sentence summary:** Tyrosine receptor signalling is tightly associated with choroid plexus enlargement and is key in CSF dynamics during a neuroinflammatory attack in MS

## INTRODUCTION

Multiple sclerosis (MS) stands as the most prevalent chronic neuroinflammatory disease, causing progressive disability. Regarding the complex and dynamic pathogenesis of MS, a growing body of evidence has posed the importance of communication between the central nervous system (CNS) and peripheral immune factors. In this particular context, the choroid plexus (ChP) has garnered significant attention as a crucial structure for the regulation and propagation of inflammation at the barrier between blood and the brain’s extracellular fluid or cerebrospinal fluid (CSF (*1*), acting as a nexus for immune cell trafficking, blood-brain barrier (BBB) regulation, cytokine production, and antigen presentation (*2, 3*). Regulating the entry of inflammatory cells into the brain and the trafficking of immune cells from brain tissue into the CSF, the ChP plays a crucial role in both maintaining neuroimmune homeostasis and responding to inflammatory and neurodegenerative brain conditions.

The choroid plexus is thought to critically contribute to the development and progression of MS (*3-5*). Recent studies found ChP enlargements in patients with MS as compared to healthy controls and demonstrated an association with functional impairment and disease progression (*6-8*). A recent study showed that a larger ChP in MS patients as compared to controls may serve as a surrogate marker for tracking neuroinflammation dependent disease progression, as well as being a marker for therapeutic responses (*6*). The latest work also highlights also the potential of ChP as a translational brain imaging biomarker for the quantification of neuroinflammation in humans and mice.

In recent years, transcriptomic approaches have been established as powerful tools for unravelling molecular pathways and mechanistic networks associated with disease initiation and progression in various neuroinflammatory and neurodegenerative disorders. Moreover, CSF proteomics were shown to have the potential to contribute to the development of diagnostic tests, prognostic markers, and therapeutic targets for various neurological disorders. Changes in specific proteins or protein patterns provide illustrative insights into disease mechanisms and might aid in early disease stages to dissect ongoing pathological abnormalities. In MS, neurofilament light chains (NFL) analyses were specifically linked to acute injuries during relapses, as well as to neurodegeneration and the extent of axonal damage (*9, 10*). Moreover, a number of proteins related to inflammation in the CSF have been associated with the duration and advancement of MS (*11, 12*). Building upon these insights, our study endeavors to explore the association between MRI-derived volumetric measures of the choroid plexus (ChP) and CSF proteins implicated in neurobiological processes and neurological disorders. We aim to elucidate the significance of ChP-related neuroinflammation by integrating translational insights from transcriptomic data obtained from the experimental autoimmune encephalomyelitis (EAE) mouse model. By investigating the interplay between ChP volumetrics and CSF protein profiles, our research seeks to contribute to a deeper understanding of neuroinflammatory mechanisms and potentially unveil novel avenues for diagnosing and managing neurological disorders.

## RESULTS

### Clinical characteristics

In our study, MRI, clinical and CSF proteomic data from 69 RRMS patients (discovery cohort) and 30 RRMS patients (replication cohort) were included. All patients were included during a relapse period, thus, depicting acute neuroinflammatory damage. There was no significant difference between the discovery cohort and the replication cohort on the main clinical characteristics (EDSS, disease duration, mean age at CSF sampling and MRI). The demographic and clinical data of the MS patients are summarized in table 1 after separating the MS patients in the discovery cohort from the replication cohort. The comparison of ChP volumes with total brain volume (TBV) normalization between the discovery and replication cohorts showed no significant difference just as the comparison of ventricle volume between both of the cohorts (Figure 1). However, both cohorts showed an increased ChP volume in comparison to the healthy controls (HC) (ANOVA F_(131,2)_=7.98, p=0.0006).

**Table 1.** Clinical data of the discovery cohort and replication cohort.

| Clinical data | HC<br>(n = 35) | Discovery<br>cohort<br>(n=69) | Replication<br>cohort<br>(n= 30) | Test statistic | P-value |
| --- | --- | --- | --- | --- | --- |
| Sex (female[%]) | 23 [64] | 38 [55] | 22 [73] | 2.4* | 0.12 |
| Mean age ± SD<br>years | 30.2±9 | 31.8±8.5 | 32.4±9.2 | 0.75<sup>†</sup> | 0.48 |
| Median EDSS ±<br>SD |  | 2±0.8 | 2±1.1 | 2.24<sup>†</sup> | 0.03 |
| Disease duration in<br>months as mean<br>(range) |  | 0 (0-4) | 0 (0-3) | 1.3<sup>†</sup> | 0.21 |

\*group comparisons performed using a $\chi^2$ test
<sup>†</sup> group comparisons performed using ANOVA
HC = healthy controls; SD = standard deviation; EDSS = Expanded Disability Status Scale.

**Fig. 1.**
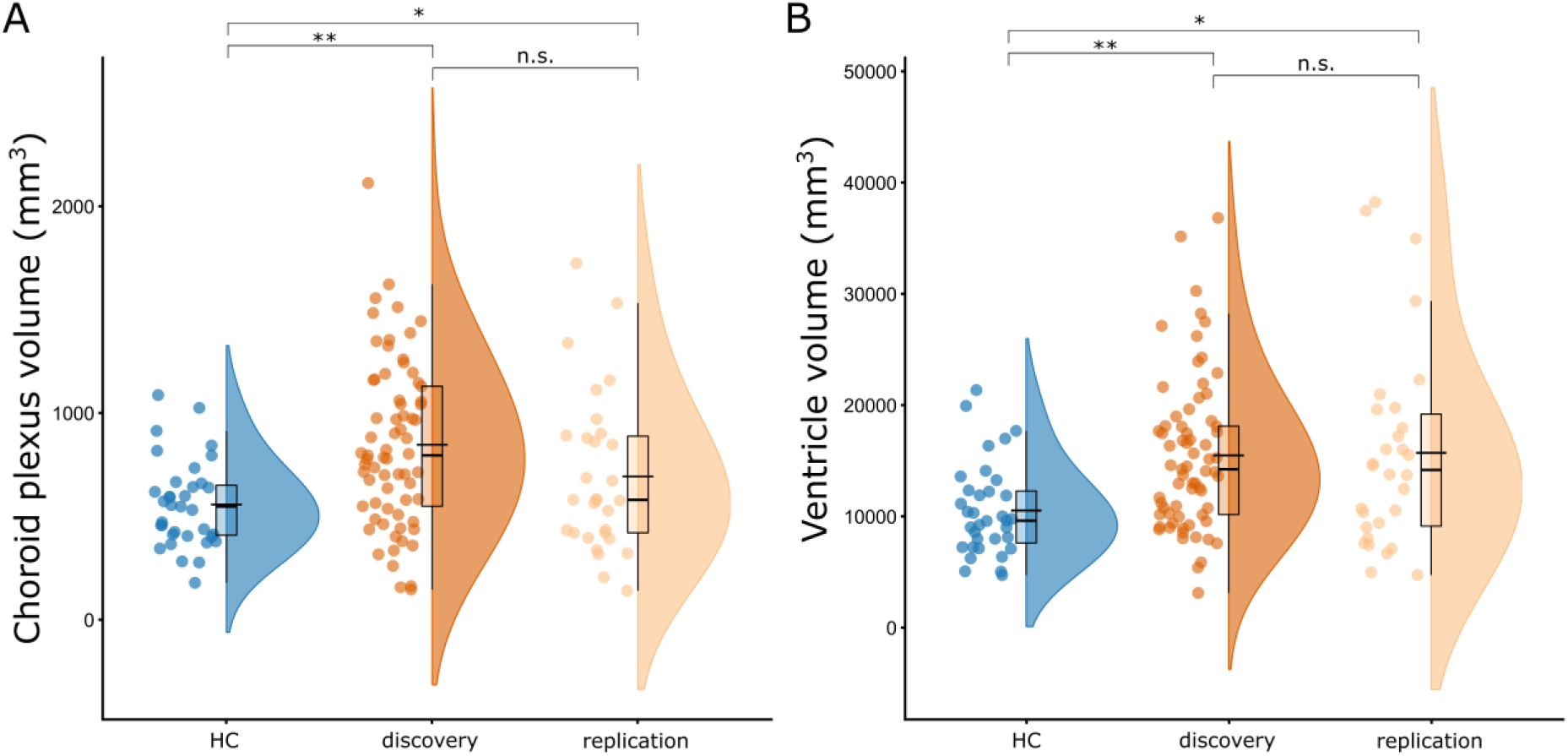
Comparison of ChP volumes between the healthy controls (HC; blue), discovery (orange) and replication cohorts (light orange; left panel) and ventricle volume between the HC (blue), discovery (orange) and replication cohorts (light orange). Age, sex and total brain volume (TBV) were used as covariates in the model. *p<0.05, **p<0.001, n.s. (not significant).

#### Associations between ChP integrity and CSF proteins spectra

In the discovery cohort, 25 of the 92 proteins analysed were significantly associated with ChP volume. In the replication cohort, ChP enlargement was linked with 10 proteins, with five of the proteins being identical to those identified in the discovery cohort (Figure 2A). The depicted proteins included Neurotrophic Receptor Tyrosine Kinase 2 (NTRK2) (Table 2, discovery pFDR=0.013, replication pFDR=0.048), Disintegrin and Metalloproteinase Domain containing Protein 23 (ADAM23) (Table 2, discovery pFDR=0.019, replication pFDR=0.045), Scavenger Receptor class B, member 2 (SCARB2) (Table 2, discovery pFDR=0.028, replication pFDR=0.05), Carboxypeptidase M (CPM) (Table 2, discovery pFDR=0.027, replication pFDR=0.044), and Contactin-5 (CNTN5) (Table 2, discovery pFDR=0.033, replication pFDR=0.041).

**Table 2.** Proteins associated with choroid plexus volume in the discovery and replication cohorts.

|  | Discovery cohort |  | Replication cohort |  |
| --- | --- | --- | --- | --- |
|  | r | P* | r | P* |
| NTRK2 | 0.382 | 0.013 | 0.44 | 0.048 |
| ADAM23 | 0.37 | 0.019 | 0.445 | 0.045 |
| SCARB2 | 0.356 | 0.028 | 0.392 | 0.05 |
| CPM | 0.334 | 0.027 | 0.4 | 0.044 |
| CNTN5 | 0.327 | 0.033 | 0.451 | 0.041 |

\*Presented P-values are corrected for multiple comparisons using FDR. r = r-score.

**Fig. 2.**
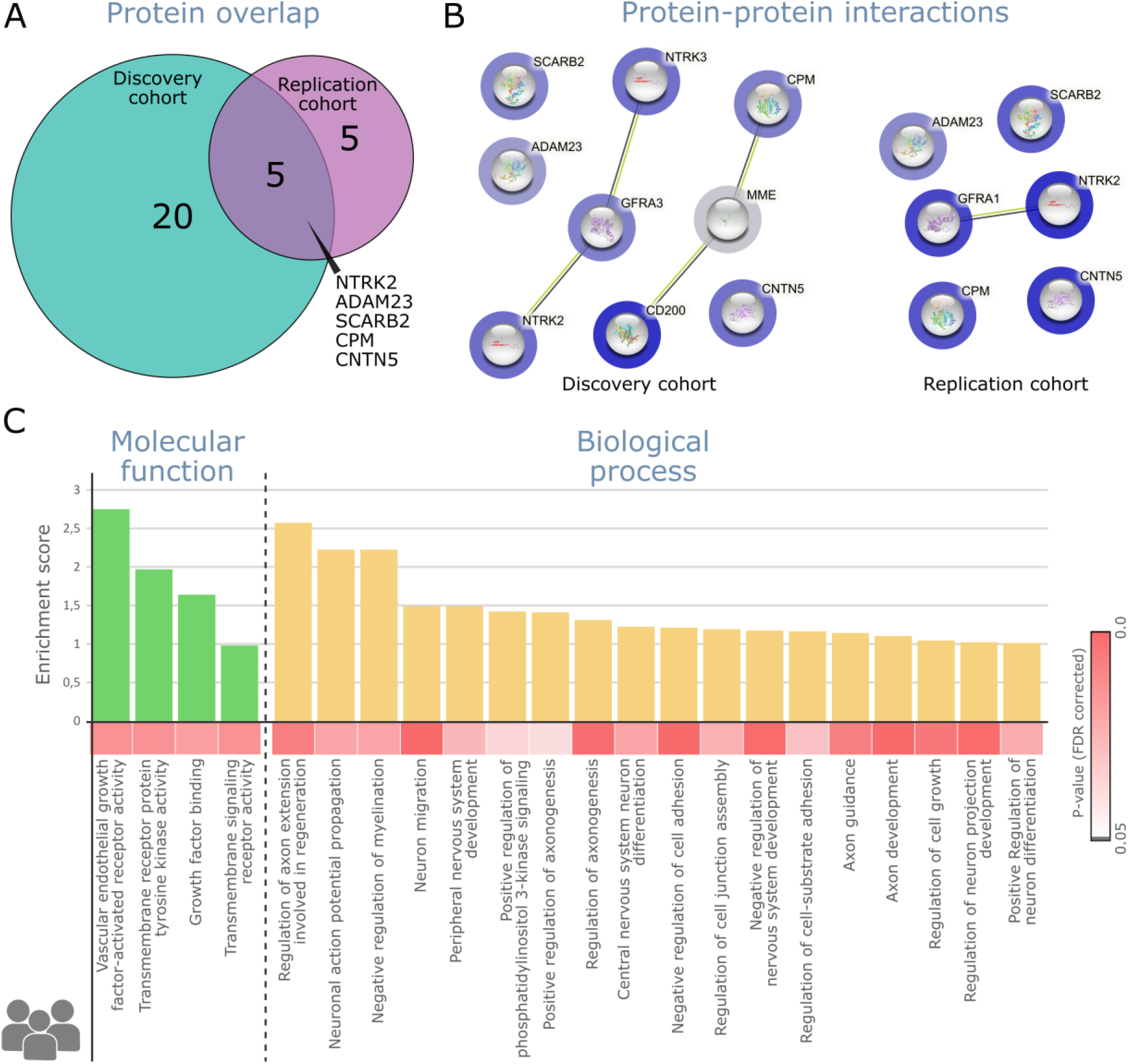
Protein–protein interactions and functional annotation analyses. A) Venn diagram illustrating the proteins associating with the choroid plexus volume in both, the discovery (green) and replication (light purple) cohorts, as well as the overlapping proteins. B) Protein-Protein interaction (PPI) network for both the discovery (left) and the replication (right) cohorts. The blue scale colours on the circles indicate how strongly represented the proteins are in each sample, where darker colors indicate highly represented proteins. C) Downstream analyses. Significant enrichment for molecular function (green) and biological process (yellow) pathways.

#### Protein enriched network analysis

The PPI network, depicted in Figure 2B and constructed from proteins associated with ChP volume, demonstrated notable similarities in both the discovery cohort (enrichment pFDR = 0.0042) and the replication cohort (pFDR = 0.046) in terms of the most enriched genes. The identified network displayed enrichment in 22 overlapping pathways, of which four were related to molecular functions and 18 were related to biological processes (Figure 2C). The enriched molecular function pathways were mechanisms of cellular communication (i.e., vascular endothelial growth, transmembrane tyrosine receptor activity, and signalling), as well as growth factor receptor activity. Regarding the biological process, the significant enriched functional pathways were related to regulation of plasticity (i.e., axon modifications and myelin regulation), as well as to neuronal regulation (i.e., neuron generation, differentiation, and migration).

#### EAE ChP proteomics analysis

To corroborate the findings obtained from the CSF of MS patients, we conducted a comparative analysis with the transcriptomic profile derived from the ChP tissue specific to EAE. The ChP samples extracted from the mice evidenced the involvement of immune molecule activity (particularly cytokines) and cellular communication (also involving receptor signalling activity and transmembrane receptors) as molecular processes; and immune and inflammation related pathways (involving leukocytes and lymphocytes) as biological processes (Figure 3A; upper panels). A detailed overview of the identified proteins within these enriched pathways is provided in supplementary table 1. The significant participation of tyrosine kinase proteins, particularly NTRK2, in influencing ChP volume within the human cohort compelled us to prioritize investigations into tyrosine-related pathways. The targeted search analyses using terms involving ‘tyrosine’ revealed five related pathways (Figure 3B) implicated in signalling and activity of the tyrosine kinase protein. The analysis of the ‘vascular’ term revealed four enriched pathways (Figure 3B), comprehending the regulation of vasculature and the adhesion of leukocytes to vascular endothelium, which is a hallmark of inflammatory processes. However, the investigation of the ‘endothelium’ term alone did not result in any significant enrichment.

**Fig. 3.**
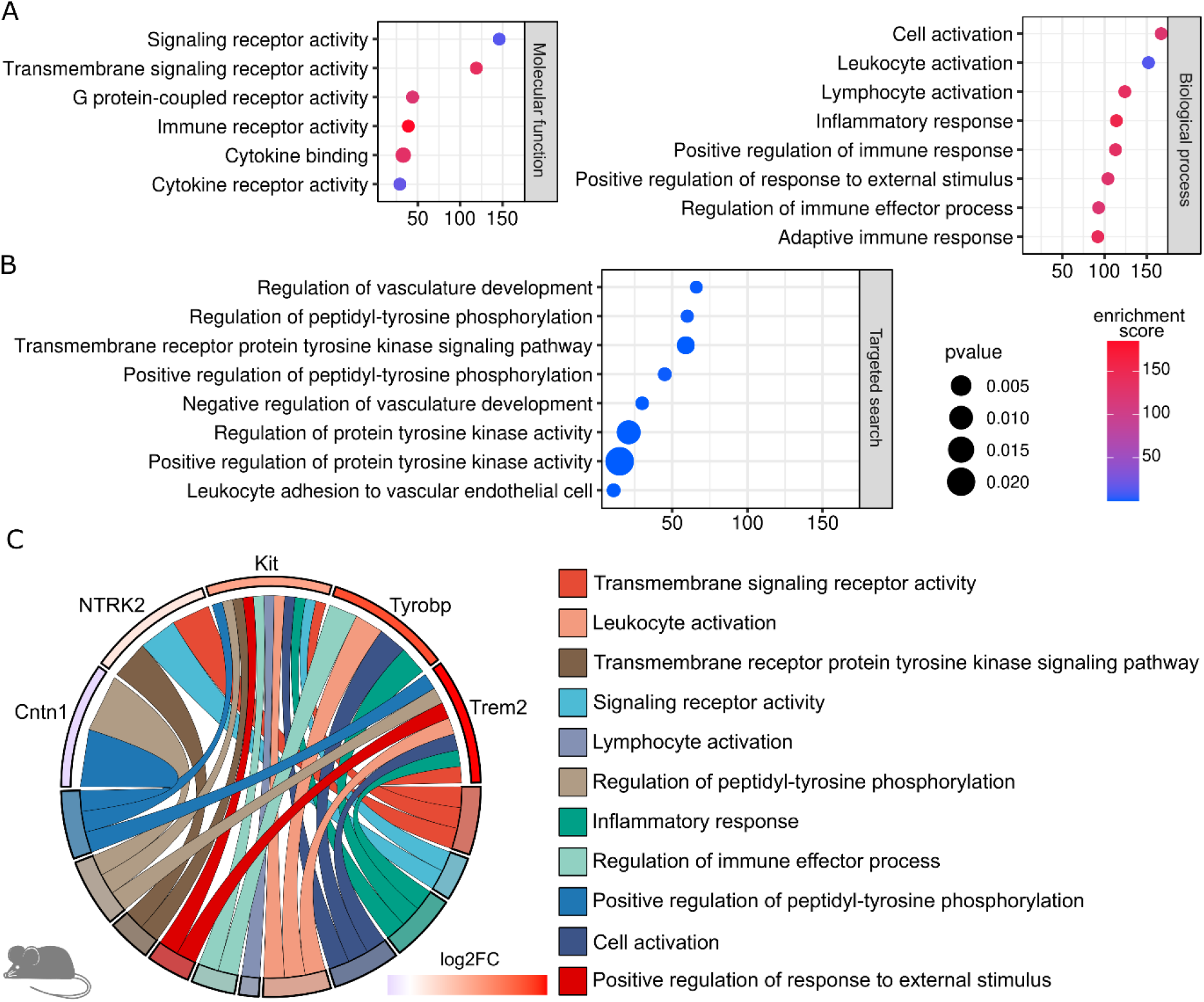
Protein-interaction analyses in the mice model of inflammation. For these analyses, experimental autoimmune encephalomyelitis (EAE) mice were compared at the peak of inflammation (14 days) against baseline expression. The resulting fold2log change was used to create a Protein-Protein interaction (PPI) network depicting pathways that are differentially expressed during neuroinflammation. The downstream analyses evidenced significant enrichment of pathways similar to those in the human cerebrospinal fluid (CSF) samples (A). Additionally, the targeted analyses depicted further tyrosine kinase enriched pathways (B). C) The pathway chord diagram depicted that genes from the tyrosine protein kinase family (Cntn1, NTRK2, Kit, and Tyrobp) as well as tyrosine receptors (Trem2) were the most abundant among enriched pathways. In A and B, X-axis reflects the gene counts.

#### Genetic variation and gene expression across reference human brain samples depicts local susceptibility to NTRK2

Our current analyses have highlighted NTRK2 as a significant translational marker in all enriched pathways for humans and mice (Fig 2 A and Fig 3 C). Next, the GTEx analysis revealed that the basal ganglia, especially the putamen, exhibited the highest levels of NTRK2 expression, followed by the frontal cerebral cortex (Figure 4A). Sex-biased expression in the included regions was tested for using bulk gene expression. However, despite the putative higher expression observed in female donors, there were no sex differences observed in the regions of the basal ganglia or the cerebral cortex (Figure 4B).

**Fig. 4.**
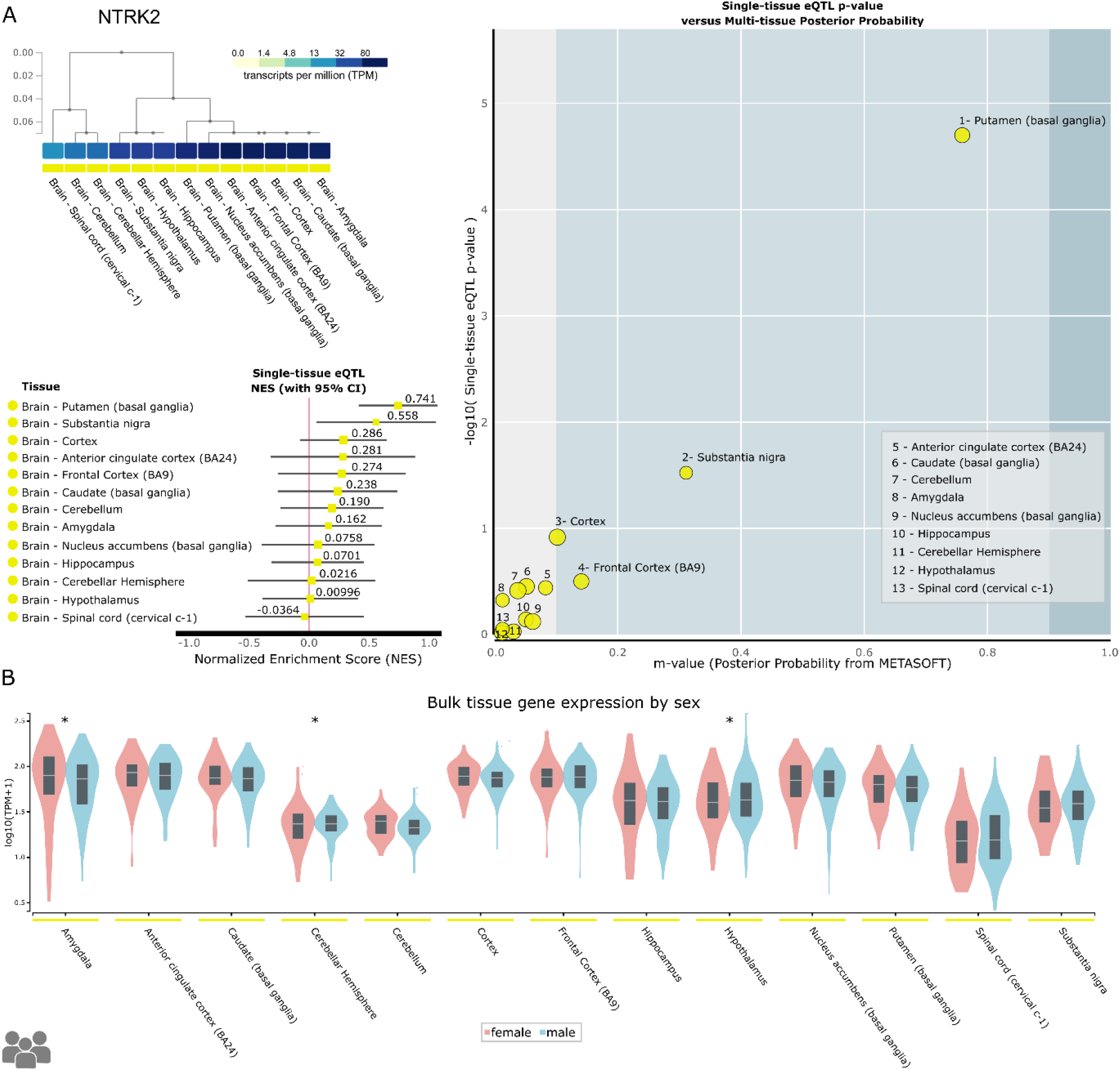
Results of the GTEx analysis for NTRK2. A) Regional susceptibility analyses showing the basal ganglia (putamen) and cortex (frontal area) as the regions with the highest expression of NTRK2 gene. B) Bulk gene expression for all regions depicted no sex-bias in the expression of NTRK2 gene across the basal ganglia or cortical regions. Asterisks depict significant differences between male and female.

## DISCUSSION

In this study, we assessed the involvement of the ChP in neuroinflammation during relapses in MS patients through proteome profiling of human CSF on a molecular basis. We further conducted RNA sequencing of ChP tissue in the EAE mouse model to mechanistically investigate the related patterns of neuroinflammatory attacks and the similarities in humans and mice. We identified five highly enriched proteins associated with ChP hypertrophy during relapses in both studied MS patient cohorts. Of the five proteins examined, NTRK2 emerged as a reliable translational indicator of acute neuroinflammation in both humans and mice. Our extensive translational analyses showed a significant involvement of tyrosine kinase pathway signalling during the inflammatory phase in MS. This mechanism may orchestrate cellular responses and tissue damage through cellular signalling and maintenance of the inflammatory response. Two other proteins, ADAM23 and CPM, were related to cell adhesion through interactions with integrins and other adhesion molecules as well as kinin-kallikrein system activation in neuroinflammation.

In a recent study, we demonstrated that the enlargement of the ChP in MS aligns with neuroinflammatory processes emphasizing its pivotal role in mediating interactions between the peripheral and central immune systems (*6*). Together with the BBB, the blood CSF barrier (BCSFB) in the ChP acts as a selective gatekeeper for immunoinflammatory response in the CNS. Despite these insights, the biological mechanisms underlying volumetric alterations in the ChP in MS patients have remained elusive.

Post mortem studies in MS patients have unveiled an accumulation of antigen-presenting cells in the ChP stroma, disruption of tight junctions within the ChP epithelium, and an activation of immune cells (*5, 13*). Under homeostatic conditions, the endothelium of the ChP is physiologically permeable (in contrast to the BBB). Moreover, peripheral inflammatory processes like intestinal inflammation may influence ChP permeability (*1, 14*). However, the molecular mechanisms underlying these dynamic ChP abnormalities were not previously studied. Therefore, our study offers a first mechanistic perspective of the processes pertaining to the ChP integrity, as measured with MRI and CSF dynamics during inflammation.

To gain a better understanding of the pathophysiological mechanisms dependent on the ChP during in an inflammatory attack, we have dissected the biological and molecular pathways in which the observed CSF proteins operate. The identified pathways encompassed cellular communication and signalling (such as tyrosine kinase receptor activity and binding) as well as cell migration and plasticity induction (including cellular generation, differentiation, and migration). Consistent with this, a recent study by Elkjaer et al., (*11*) reported “cellular migration” as a major characteristic enriched in all MS subtypes, including relapse, secondary progressive, and primary progressive MS. Our study depicts a more in-depth view with the description of the transmembrane receptor protein tyrosine kinase activity followed by the transmembrane signalling and vascular endothelial growth factor receptor activity as key mechanisms in the molecular functional pathways analysis. The key pathways detected indicate the involvement of inflammatory and vascular endothelial functions as key modifiers in ChP hypertrophy for MS patients in relation to acute neuroinflammation. Another vital point is the detection of these pathways in the conducted EAE mouse model. Recognizing the presence and relevance of these pathways in an in vivo model that closely recapitulates human disease conditions enhances the likelihood that our findings could be translated into meaningful therapeutic strategies for MS patients.

Regarding the central role of NTRK2, protein tyrosine kinases, which are the most important factors found in our study, are key components of various signalling pathways that regulate immune cells, including T and B lymphocytes. In MS, tyrosine kinases directly modulate the functions of B cells, macrophages, and microglia therefore targeting both adaptive and innate mechanisms that contribute to the immunopathology of MS on both sides of the BBB (*15*). Activated tyrosine kinases downstream of immune cell receptors contribute to immune response promotion, inflammation, and tissue damage. Given the critical involvement of tyrosine kinases in MS pathology, targeted therapies aimed at modulating immune responses may help better mitigate the disease’s impact on patients’ quality of life. An example of this is the treatment with CNS-penetrant BTK (Bruton tyrosine kinase) inhibitors. This treatment option is currently under efficacy evaluation in clinical trials, but preclinical studies in the EAE model have shown promising results in which key pathological features of MS, including B cell activation, CNS lymphocyte infiltration, leptomeningeal inflammation, pro-inflammatory microglial activation, and demyelination can be supressed (*16*). See Kramer et al., (*15*) for a recent overview on this topic.

Regarding specific attested proteins, the tyrosine kinase NTRK2 showed a robust correlation with ChP volumes in both cohorts and was the highest enriched gene on both humans and mice. NTRK2 plays an essential role in biological pathways including brain plasticity, impacting neuron survival, proliferation, migration, differentiation, and synapse formation. All these pathways were indeed enriched in our study. NTRK2 is a specific receptor of the Brain-Derived Neurotrophic Factor (BDNF). The binding results in the activation of the MAPK pathway and regulation of synaptic plasticity and repair (*17*). Furthermore, it can bind NTF4/neurotrophin-4 and NTF3/neurotrophin-3, which regulates neuron survival. A recent genetic study, detected that the NTRK2 gene (of note, together with the STAT3 gene) has a significant overlap in the genetic susceptibility of MS and linked psychiatric comorbidity (*18*). In their paper, the NTRK2 gene had the highest enrichment and participated in the main signalling pathways (i.e. immune interaction, cytokine responses) that were identified. In line with these findings, in our study, NTRK2 showed high PPIs and involvement in the enriched pathways in both cohorts. Overall, the involvement of NTRK2 in the ChP of MS patients suggests its potential significance in regulating neuroinflammatory processes, maintaining barrier integrity, and providing neurotrophic support.

The GTEx data revealed the highest expression of NTRK2 in the basal ganglia, particularly in the putamen, of the human brain. The involvement of the putamen in MS has been established in untreated patients with clinically isolated syndrome (CIS) (*19*) and in patients with different types of MS (*19-21*), with increased vulnerability to lesion formation and demyelination (*22*).

The examined mouse model has highlighted immune-related and cellular communication pathways that closely mirror the human results. This underscores the sensitivity of CSF measurements to detect molecular abnormalities in ChP tissue. Additionally, the targeted search analyses reiterated a significant involvement of tyrosine kinase metabolism and vascular components in neuroinflammation in both humans and mice. Given, that the ChP integrity is associated with both inflammatory activity and specific proteins, inhibition of the function of these molecules or their expression may influence disease activity. Thus, our findings suggest the potential for ChP as a therapeutic target whilst highlighting its mechanistic implications in neuroinflammation and neurodegeneration.

This study has limitations. The different MRI acquisition protocols in the two patient cohorts may introduce variations in ChP segmentation. However, there were no significant differences in the resulting volumes between both cohorts. Moreover, both groups of patients were recruited with the exact same clinical inclusion criteria and protein sequencing, therefore, reducing the bias on clinical and sampling acquisitions settings. Nonetheless, further validation in larger cohorts is essential to confirm the detected non-inflammatory pathways and compensatory effects amid ChP characteristics.

Overall, our results provide new insights into the underlying mechanisms of acute neuroinflammation in MS through non-invasive ChP characterisation, protein expressions in CSF, and the description of the underlying mechanisms on a mesoscopic and molecular basis. Given the complexity and dynamic pathogenesis of MS, our results highlight the possible future role of tyrosine kinase pathway modulation targeting immune cell activation and migration. Depending on the specific targets, strong immunomodulatory effects could potentially impact the inflammatory processes seen in MS. Overall, our research offers novel prospects to thoroughly investigate multiscale networks and mechanisms within the inflammatory CNS attacks, with a focus on the interplay between the peripheral and central immune system. This could pave the way for innovative therapeutic interventions that might modify the disease trajectory of MS.

## MATERIAL AND METHODS

### Ethics statement

The study was conducted in accordance with the Declaration of Helsinki and approved by the local ethics committee (ethics committee of the Landesärztekammer Rhineland Palatinate, number 837.479.17 [discovery cohort] and number 837.019.10 [replication cohort]). Written informed consent was obtained from all participants.

### Patient cohorts

Patients form the discovery cohort belonged to larger MS cohort (n = 1156) with prospective comprehensive and standardized clinical and standardized 3T MRI data collection from the Department of Neurology at the University Centre Mainz. For the current analyses a group of 69 patients with relapsing remitting multiple sclerosis (RRMS), confirmed according to the revised 2010 McDonald diagnostic criteria, cursing a relapse period, and having CSF sampling for protein analyses were included. The replication cohort consisted of patients from the outpatient clinic, which were diagnosed using the same clinical criteria and but their MRI protocol was not standardized.

For the discovery cohort, individual images were acquired on 3T MRI scanner (Siemens Skyra) with a 3D T1 MPRAGE axial sequence with following parameters: Echo Time (ET) = 0.0025, Repetition Time (RT) = 1.6, Inversion Time (IT) = 0.9, Flip Angle (FA) = 8°, matrix size = 200x256x192, voxel size = 0.9mm^3^. The second (replication) cohort (n=30) was included from the same department. For the replication cohort, the scans were acquired on the same scanner using a modified 3D T1 MPRAGE sagittal sequence with following parameters: ET = 0.0024, RT = 1.9, IT = 0.9, FA = 9°, matrix size = 192x256x256, voxel size = 1mm^3^. Each patient was clinically assessed by an experienced neurologist, and the Expanded Disability Status Score (EDSS) score was determined at disease onset (study entrance), annually for two years, and after four years. In total, MRI, clinical, and CSF proteomic data from 99 RRMS patients were available and were included in the study.

### Human MRI processing

Magnetic resonance imaging preprocessing was performed using the open-source FreeSurfer (https://surfer.nmr.mgh.harvard.edu/) software, which is currently the most widely used software to automatically segment the brain structures, including the ChP. Details on the segmentation procedures can be found elsewhere (*23*). In brief, FreeSurfer uses a probabilistic atlas built from manual segmentations of a training dataset normalized to the MNI305 space. This allows a point-to-point correspondence between all the training subjects. The atlas provides the probability of each label at each voxel, the probability of each label given the classification of neighboring voxels (neighborhood function), and the probability distribution function of voxel intensities, modelled as a normal distribution, for each label at each voxel. The segmentation of a new individual MRI is achieved by spatially registering the new subject to the MNI305 space and incorporating the subject-specific voxel intensities to find the optimal segmentation that maximizes the probability of observing the input data. FreeSurfer allows the segmentation of both the ChP and the CSF. Segmentation using FreeSurfer was visually inspected and corrected where necessary for each subject.

### Human protein analyses

CSF sample collection was performed via lumbar puncture according to standard procedures (*24, 25*). The CSF samples were then assayed using the proximity extension assay technology (PEA) (Olink Proteomics AB, Uppsala, Sweden), in which 92 oligonucleotide-labelled antibody probe pairs representing proteins related to the nervous system are allowed to bind to their respective target present in the sample (Olink Target 96 Neurology panel). The PEA technique has a high specificity and sensitivity (*26*). The platform provides Normalized Protein eXpression (NPX) data where a high protein value corresponds to a high protein concentration, but not to an absolute quantification. NPX are obtained by a series of computations. These operations are designed to minimize technical variation and improve interpretability of the results. All assay validation data (detection limits, intra- and inter-assay precision data, etc.) are available on the manufacturer’s website (www.olink.com).

In order to investigate the biological activities that are regulated through the identified ChP associated proteins, we explored protein–protein interactions (PPIs) using the STRING database (*27*), by using the correlation values as ranks for the ontology and enrichment (*28, 29*). For functional annotation of the significantly associating proteins, we effectuated Gene Ontology analysis within STRING. This allowed for the identification of specific biological and molecular pathways to which these significantly different proteins belong to.

To establish a link between the observed pathways and inherited brain susceptibility to disease, we employed the Genotype-Tissue Expression (GTEx) portal (https://gtexportal.org/home/) by searching the gene(s) that were commonly enriched in both humans and mice (see more details on mouse experiments below). Correlations between genotype and tissue-specific gene expression levels help identify regions of the genome that influence whether and how strongly a gene is expressed. GTEx helps researchers to understand inherited susceptibility to disease. The dataset included 205 samples, of which 156 corresponded to male donors. This allowed for examining, using bulk gene expression, of sex-bias in all available brain structures.

### EAE mouse experiments

EAE (experimental autoimmune encephalomyelitis) was induced in C57BL6J mice (N = 5; females, 9 weeks old at the start of treatment, Envigo) using a subcutaneous injection of 200mg of MOG peptide (Myelin Oligodendrocyte Glycoprotein Peptide Fragment 35 to 55; from Charité) emulsified in complete Freund’s adjuvant (from Sigma-Aldrich) that contained 200mg of Mycobacterium tuberculosis H37RA (from Difco). Pertussis toxin (400 ng; Enzo Life Sciences) was injected intraperitoneally in 200mL phosphate-buffered saline (PBS) on the day of immunisation and again two days later. Two independent investigators scored disease severity daily in an anonymised manner using a 0 to 5 scale (EAE score), as described elsewhere (*30*). The peak of demyelination occurs 10 to 15 days after injection.

#### RNA sequencing

Choroid plexus (ChP) tissue was obtained for bulk RNA sequencing analyses from both naive and EAE (day 14) samples. Using an established protocol (*6, 31*), ChP tissue was manually dissected from the lateral, third, and fourth ventricles with the aid of an illuminated stereo microscope. The procedure involved digestion of tissue from a single mouse in 300 µL of Hank’s balanced salt solution (HBSS) (Gibco; Catalogue number: 14025-092) containing collagenase and dispase (Merck; Catalogue number: 11097113001; concentration: 0.1 mg/mL) for 30 minutes at 37 °C using an orbital shaker. The homogenised tissue was then passed through a 70-µm pore size cell strainer using an insulin syringe and washed with 600 µL HBSS solution. Finally, the sample was centrifuged at 500 × g for 5 minutes at room temperature. The supernatant was discarded, and the cellular pellet was suspended in 350 µL of RLT buffer. RNA isolation was carried out by using a Qiagen RNeasy Micro Kit (Catalog No. 74004) in accordance with the instructions provided by the manufacturer. The quality and quantity of RNA were confirmed using NanoDrop and Bioanalyzer RNA 6,000 nano Kit (Agilent). Samples with RNA integrity number values exceeding 6.5 were used for RNA sequencing. NEBNext ribosomal RNA depletion was conducted, followed by NEBNext directional Ultra RNA II Library preparation and sequencing on the NextSeq500 platform (Illumina) using the high output version 2 kit with 75 cycles. To eliminate low-quality reads, fastp (*32*) was employed, and overrepresented sequences were analyzed. In addition, polyG and polyX tail trimming was performed with parameters set to -g -x -p. The data quality of the trimmed output was verified using fastqc. The data were aligned to the most recent reference genome for mice (GRCm39) using the Spliced Transcript Alignment to a Reference aligner, which is designed for long-reads. Samtools filtered out low-quality alignments, leaving only high-quality alignments that were quantified using StringTie. The org.Hs.eg.db and org.Mm.eg.db databases were used for annotation.

Next, we analysed the variance in gene expression between naive and EAE (14-day) mice via the DESeq2 package on R-studio. This was followed by a protein-related gene search to assess protein-interaction networks. Specifically, we utilized the Search Tool for the Retrieval of Interacting Genes (STRING) database (http://www.string-db.org/) with a confidence score of 1 to form the foundation for the functional study of the proteome. To examine the translational properties of the discovered networks in the human proteome, we conducted a targeted search on the EAE data. To do so, we utilized PiNGO (Version 1.5.2; http://www.psb.ugent.be/esb/PiNGO) in Cytoscape 3.9.1, a tool that identifies candidate genes in biological networks that are predicted to be involved in specific processes or pathways of interest. In summary, PiNGO permits a limited exploration of the ChP protein coding genes’ engagement in specific acknowledged roles via a straightforward network-based technique and Gene Ontology categorization systems. PiNGO evaluates enrichment statistics using a hypergeometric test and regulates the resulting P-values for several analyses using Benjamini-Hochberg FDR corrections (*33*). We conducted the target search utilizing terms linked to ‘tyrosine’, ‘vascular’, and ‘endothelium’ pathways.

### Statistics

Relevant proteins spectra were initially identified by spatially correlating NPX protein levels with ChP volumes. Nuisance variables such as the total brain volume, age, and sex were used. Correction for multiple comparisons was completed at a false discovery rate (FDR) of 5%, indicated by “pFDR”. To ensure the robustness of the associations, the same set of analyses was performed in the replication cohort.

## Acknowledgements

The authors thank all the patients who participated in the study and Kathleen Claussen for proofreading this manuscript.

## Author contributions

Conceptualization: G.G-E., S.G., F.L.

Methodology: G.G-E., T.R., J.G., P.S.W.

Investigation: G.G-E., J.S., F.L., S.G.

Visualization: G.G-E.

Funding acquisition: S.G., G.G-E., F.Z., O.S., F.L., V.F.

Project administration: F.L., V.F., J.S., P.S.W., J.G., T.B., S.G.M.

Supervision: S.G., F.L.

Writing – original draft: G.G-E., J.S., F.L.

Writing – review & editing: F.L., J.S., G-G-E., V.F., S.E., D.C., T.K., P.S.W., J.G., T.B., A.O., S.B., S.G.M., F.Z., O.S., S.G.

## Data and materials availability

All the current work was performed using openly available software: https://surfer.nmr.mgh.harvard.edu/, http://www.string-db.org/, http://www.psb.ugent.be/esb/PiNGO, https://gtexportal.org/home/. Data cannot be made openly available due to institutional restrictions, but it can be made available from the corresponding authors upon reasonable request and the corresponding data transfer agreement (DTA).

